# Doxycycline Release From Cyclodextrin Oligomer-Containing Collagen Gels

**DOI:** 10.1101/2025.05.28.656660

**Authors:** Eric Trout, Leena Palomo, Horst A. von Recum, Steven Eppell

## Abstract

Dental implants commonly suffer from chronic peri-implantitis arising from infection and inflammation at the abutment/gingiva interface, primarily because they fail to replicate the collagen-rich soft tissue interface between natural teeth and the jawbone. Current treatment strategies involve frequent administration of systemic antibiotics and collagenase inhibitors which complicates clinical management. A localized controlled drug-release approach may offer a way to simplify this clinical management. In this study, we investigate doxycycline loading and release from a collagen hydrogel containing entrapped oligomers of γ-cyclodextrin (CD). Incorporating these cyclodextrin oligomers increased the releasable amount of doxycycline by 220% and reduced its release rate fivefold. The resulting enhancement in both drug loading and release control expands the potential for further development of a collagen-coated dental implant.

## Introduction

Cementum of a native tooth connects to alveolar jaw bone through a soft tissue collagenous fiber interface. These fibers are the mechanical loading structure for mastication and serve as physical barriers preventing bacterial infiltration. Current dental implants do not recapitulate this tissue barrier. While over 97% of implants remain intact in the jaw, complications stemming from infection, inflammation, and the resulting challenges with aesthetics and functionality are common. Prevalence of peri-implant mucositis and peri-implantitis of 19-65% are reported(Schwarz et al., 2018). As a result, recent meta-analysis concludes even teeth compromised by periodontal and endodontic problems may have longevity that surpasses the average dental implant. Since the dental implant market is estimated to grow globally near 10% compounded annually until 2030, the complication rate of dental implants has serious financial and human health implications(Grand View Research, n.d.).

This infective and inflammatory etiology owes to bacterial insult followed by a host inflammatory cascade. Antibiotics and collagenase inhibitors from the tetracycline family are administered to combat the condition(Qian et al., 2020). However, systemic and frequent dosing poses clinical difficulties while surgical intervention involves trauma and cost(Cha et al., 2019). Non-surgical local controlled drug release promises improved success rates of dental implants without these downsides. With this in mind, we investigate a drug release system using two components: γ-cyclodextrin oligomers within a collagen hydrogel.

Figure 1 shows the chemical structure of γ-Cyclodextrin, an 8-member oligosaccharide ring with a hydrophobic “pocket.” Small drug molecules are trapped by the stability offered from the hydrophobic region resulting in affinity-based drug-loading. Release from individual binding sites is rapid (milliseconds). However, with sufficiently dense local pockets to “hop” around on, the net release is delayed (hours to months). Such affinity-based filling within a network of insoluble polymerized cyclodextrin prolonged drug release by a factor of four compared to pure diffusion from a linear dextran (non-affinity) control polymer(Rivera-Delgado et al., 2016). These results motivated our design of a matrix containing a network of cyclodextrin binding regions.

**Figure 1.**
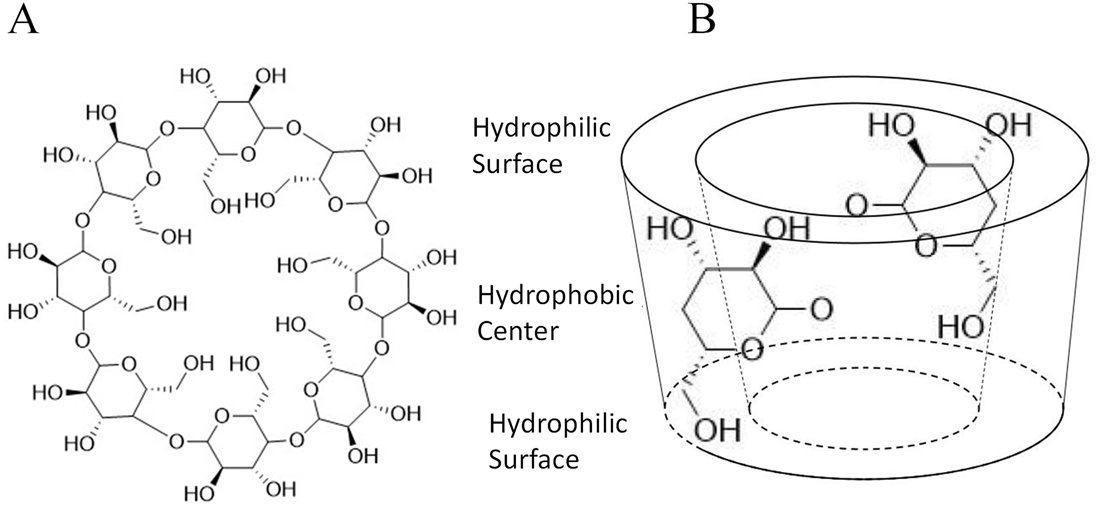
(A) Gamma-Cyclodextrin is an 8-member oligosaccharide ring. It has a moderately hydrophobic “pocket” in which drug molecules insert through affinity-based drug-loading. From the 3D model it can be seen that the hydrophilic hydroxyls form a “fringe” at the top and bottom of the ring, while the center contains the hydrophobic C-C bonds.

Collagen hydrogels may be useful matrix materials for a cyclodextrin-containing network. Type I collagen is a biocompatible protein that is easily assembled, processed, and manipulated in clinical settings. If attached to the abutment of a dental implant, collagen hydrogels could mimic the tissue seal around a native tooth.

Our process for covalently attaching collagen fibrils to titanium surfaces could provide a supracrestal fiber-like attachment of the hydrogel to the implant(Miller et al., 2020). Since diffusional distances would be short, we argue that collagen hydrogels provide an excellent platform to develop, study, and clinically translate a dental implant attached drug release system with the expectation of modest drug delivery from a thin collagen coating.

Figure 2 is a schematic of the proposed collagen matrix with cyclodextrin and doxycycline. Fibrous collagen entangles to form a hydrogel containing interfibrillar fluid filled spaces. Entrapped cyclodextrin oligomers within these spaces provide a network of binding regions. Drug molecules are either bound to cyclodextrin hydrophobic pockets, adhered to collagen fibrils, or float in interfibrillar spaces. Affinity-based drug-loading and release is dependent on both number of affinity groups and distance between neighboring groups; clusters of cyclodextrins on an oligomer perform better than dispersed cyclodextrin monomers(Cyphert et al., 2020)

**Figure 2.**
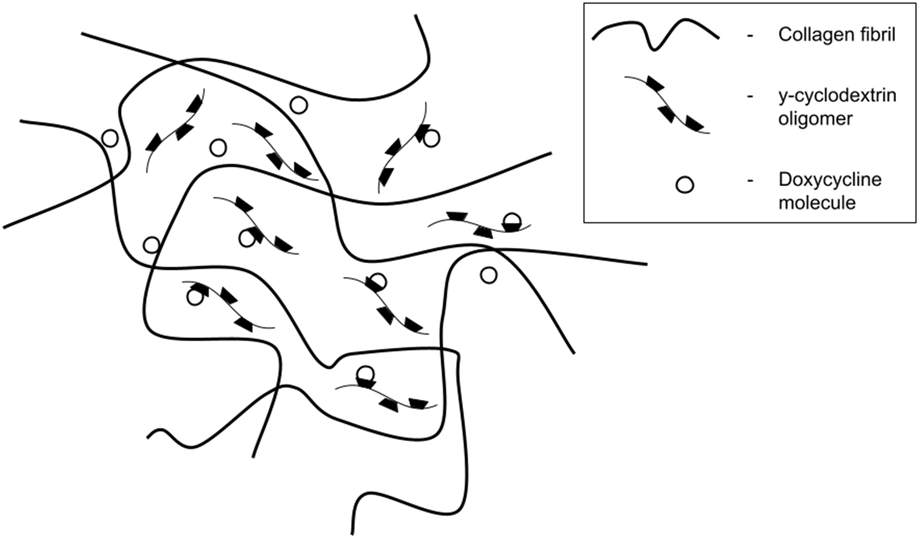
Collagen fibrils 50-100 nm in diameter entangle to form a hydrogel. Cyclodextrin oligomers are entrapped in fluid filled volumes between fibrils. Doxycycline molecules adhere to the collagen fibrils, associate with cyclodextrin hydrophobic pockets, or diffuse through interfibrillar spacings.

Similar to that work incorporating cyclodextrin into poly(methyl methacrylate), we hypothesize that the incorporation of cyclodextrin into collagen hydrogels will increase drug loading and prolong subsequent release compared to collagen hydrogels alone. We evaluate three sample groups: cyclodextrin loaded collagen hydrogels, pure collagen hydrogels, and pure polymerized cyclodextrin. We examine amount of releasable drug and rate of release. Contributions of cyclodextrin and collagen components are determined by fitting release using exponential models. The two collagen hydrogel groups are examined to compare release amount and rate. The polymerized cyclodextrin and pure collagen hydrogels are analyzed to quantify the role of their respective components within the combined system.

## Methods

### Collagen Hydrogel Preparation

Collagen hydrogels were created using the warm start method (Baskin et al., 2012). 3 ml of lightly pepsin digested calf skin type I collagen monomers at 3.1 mg/ml in 0.1 M HCl (PureCol^®^, Advanced BioMatrix, San Diego, CA) was added in a 1:5 ratio to tris buffered saline (TBS), buffered to pH 7.2 then incubated at 37°C for 24 hours. This produces gels consisting of native D-banded fibrils 50-100 nm in diameter (Eppell et al., 2018). Formed gels were removed from solution, placed on filter paper, and serially dehydrated by rinsing eight times with increasing concentrations of ethanol in DI water. Ethanol rinsed samples were loaded into a uniaxial press (Eppell et al., 2018) and a 10 kg mass applied for 24 hours to compress the sample mechanically at 3.5 MPa driving out any remaining liquid. This resulted in circular discs approximately 400 μm thick and 6 mm in diameter.

To ensure samples maintained their general shape during the drug release studies, dehydrated samples were crosslinked with a 10% w/w genipin solution in a 70-30 ethanol-water solution. A 15 μL aliquot of the genipin solution was pipetted onto each side of the sample while it was in the press then left for 10 minutes allowing the crosslinking solution to penetrate the sample. After infusion, the 10 kg mass was re-applied for 30 hours.

### Addition of Cyclodextrin to Collagen Hydrogels

To investigate the effect of adding cyclodextrin oligomers to the collagen hydrogels, samples with γ-cyclodextrin oligomers (γ-CD epichlorohydrin, Cyclo-lab, Budapest, Hungary) added to the TBS prior to addition of the type I collagen monomers were made. Powdered γCD oligomers (oligomer weight 89 kDa) were mixed into TBS solution to form a 0.025 g/ml CD/TBS solution.

### Doxycycline Loading and Release

Crosslinked dehydrated hydrogel discs were submerged in 1 ml of 1X phosphate buffered saline (PBS) containing 40 mg/ml doxycycline (Sigma-Aldrich, St. Louis, MO) at room temperature for 72 hours. Release of doxycycline was measured by placing the samples in 1.5 mL microcentrifuge tubes containing 1 mL PBS at 37°C under constant shaking and protected from light. 1 ml aliquots were removed and replace with fresh PBS hourly for the first 6 hours then every other day out to 11 days. Doxycycline concentration was determined by measuring the absorbance at 345 nm in a UV-Vis spectrophotometer (H1, Biotek, Winooski, VT)(Rivera-Delgado et al., 2016). All released doxycycline values are reported as means ± standard deviations. Spectroscopic measurements were run in duplicate. Each sample type was run three times. So, while there were 6 measurements made for each sample, we report n=3 to indicate the number of times the entire experiment was reproduced.

### Mathematical Modeling

Cumulative doxycycline release, f(t), was plotted against time. Using a commercial non-linear curve fitting app (MATLAB R2020a, Mathworks, Natick, MA), an equation of the form

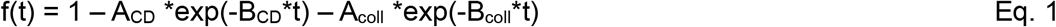

was fit to the data. Goodness of fit was determined using R-square values.

## Results

Using insoluble polymerized cyclodextrin films as a benchmark, we tested two groups of collagen hydrogels for sustained release of doxycycline: pure collagen and collagen mixed with cyclodextrin oligomers. Collagen hydrogels were loaded with doxycycline then release profiles were measured over time. Since doxycycline binds to both collagen(Albu et al., 2009; Semyari et al., 2018; Tihan et al., 2019) and cyclodextrin, (Khodaverdi et al., 2021; Rivera-Delgado et al., 2016) these two binding compartments were separated by modeling time release profiles of pure polymerized cyclodextrin and pure collagen. The B_CD_ and B_coll_ coefficients (Eq. 1) obtained from these fits were then applied to construct the model of the combined collagen/cyclodextrin system. The A_CD_ and A_coll_ coefficients (Eq. 1) were allowed to vary in the combined model to determine, quantitatively, how much doxycycline was released from each compartment.

### Cyclodextrin Incorporation

Cyclodextrin oligomers were incorporated into the collagen hydrogels non-covalently presumably in fluid volumes between collagen fibrils. No direct measurements were made to prove cyclodextrin oligomer incorporation in collagen hydrogels during fibrillogenesis; however, weighing the gels provided indirect evidence. Pure collagen gels were made using 9.3 mg of collagen for each gel. The mass of the three gels, measured after dehydration, was 8.2 mg ±0.8 mg. Cyclodextrin was incorporated using 1 ml of solution containing 25 mg of cyclodextrin. The mass of the three gels incorporating cyclodextrin, after dehydration, was 18 mg ±1.9 mg.

### Drug Loading

The hypothesis that an affinity-based mechanism of cyclodextrin(Bonfield et al., 2024; Rivera-Delgado et al., 2016) increases the amount of releasable doxycycline from a collagen hydrogel was tested by comparing final cumulative release of doxycycline in two hydrogel groups. Pure collagen samples averaged final release of 1.6 mg +/-0.22 mg, n=3. Collagen hydrogels with incorporated cyclodextrin oligomers averaged more than twice this with an average final release of 3.6 mg +/-0.30 mg, n=3. Details of these measurements are provided below.

### Controlled Long-Term Release from Collagen Hydrogels

Figure 3 shows cumulative release over time of doxycycline from collagen hydrogels loaded with cyclodextrin oligomers. Aliquots were taken every hour for 6 hours, at 24 hours and then every 2 days out to 11 days. The inset shows the first day of release in more detail. At each time point, measured release is plotted (closed circles) as a percent of the final cumulative amount released. Error bars indicate standard deviations of three separate gels fabricated and loaded with doxycycline. The plot increases rapidly during the first 8 hours then plateaus near the 12 hour mark. After the plateau, the plot slowly increases. Doxycycline release was nonzero throughout the 11 day experiment with 3.8 µg released on the eleventh day.

**Figure 3.**
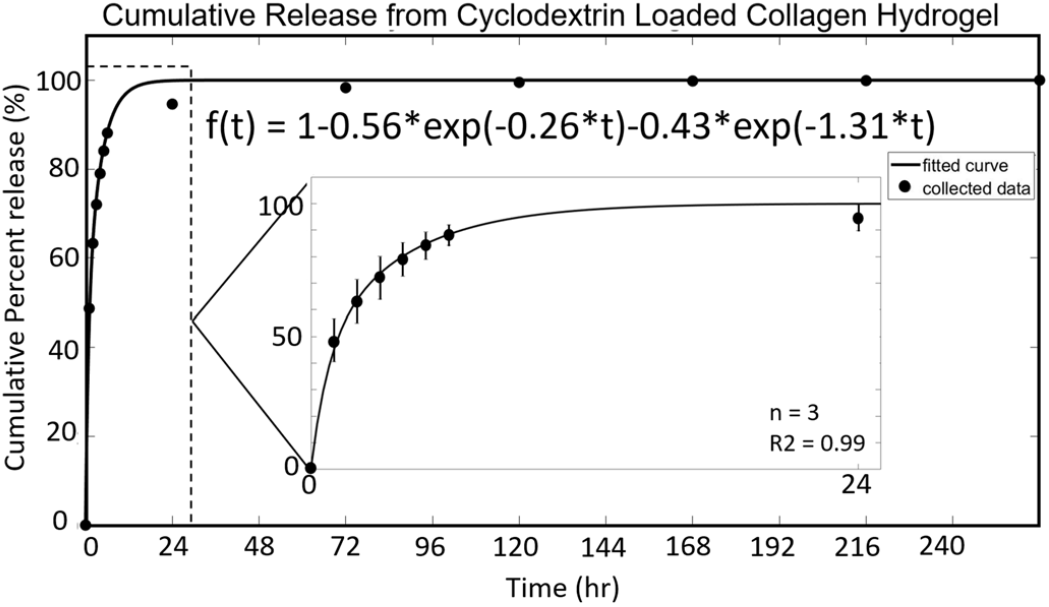
The cumulative release of doxycycline from collagen hydrogels loaded with cyclodextrin oligomers is plotted. Aliquots were taken every hour for 6 hours, at the end of the first day, and then every other day for 10 days and plotted as circles, with these time intervals being marked along the x-axis. The release amount is plotted as a percent of final cumulative amount released along the y-axis. The plot increases rapidly initially and plateaus around the 24 hour mark. After the plateau the plot increases very minimally, staying close to the asymptote of 1. A solid line is fitted to the data points. A model was created by fitting a curve in the form of y(t)=1-ACD*exp(−0.26*t)-Acoll*exp(−1.31*t). The exponent values represent the release rate from the two parts of the system: cyclodextrin and collagen, respectively. The curve fitting was performed to report back the coefficient values ACD and Acoll. These are used to provide insights into how much of the drug release the two parts of the system are responsible for. The fitted equation was 1-0.56*exp(−0.26*t)-0.43*exp(−1.31*t).

*Figure 4* shows cumulative release of doxycycline from a pure collagen hydrogel. The inset shows a quickly rising portion that plateaus between 5 and 6 hours. After the plateau, release continues slowly similar to the cyclodextrin loaded hydrogel shown in Figure 3. The 1.31 factor multiplying time in the exponent in Figure 3 was obtained from the fit shown in Figure 4.

**Figure 4.**
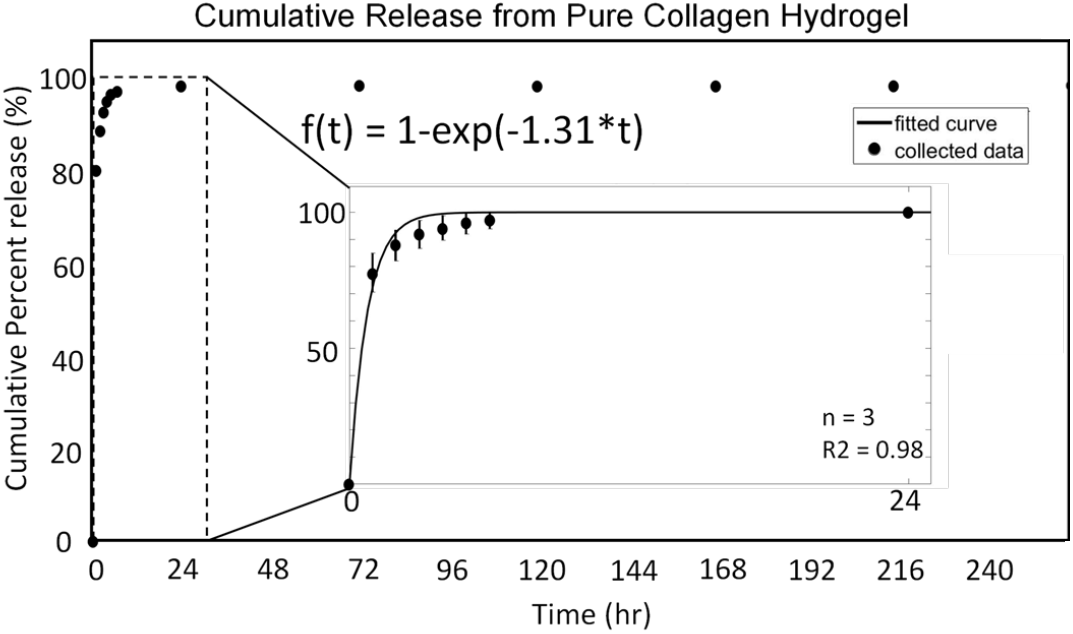
The cumulative release of doxycycline from a collagen is plotted. Aliquots were taken every hour for 6 hours, at the end of the first day, and then every other day for 10 days and plotted as circles, with these time intervals being marked along the x-axis. The release amount is plotted as a percent of final cumulative amount released along the y-axis. The plot increases rapidly initially and plateaus around the 24 hour mark. After the plateau the plot increases very minimally, staying close to the asymptote of 1. A solid line is fitted to the data points. A model was created by fitting a curve in the form of y(t)=1-exp(Bcoll*t). The fitted equation is 1-exp(−1.31*t).

### Controlled Long-Term Release from Polymerized Cyclodextrin

*Figure 5* shows cumulative release of doxycycline from polymerized cyclodextrin as reported by Rivera-Delgado et al.(Rivera-Delgado et al., 2016). That study was only conducted for 48 hours. For ease of comparison, the data plotted here runs out to 240 hours as in figures 3 and 4. There is a rapid release over about 8 hours with a plateau around 24 hours. While the 240 hour plot looks qualitatively similar to figures 3 & 4, the 24 hour inset shows a decidedly different release vs. time profile. The last 5 data points follow a smooth inverse exponential path similar to previous cases. However, earlier data points are arrayed in roughly linear fashion offset below the exponential path by a few tenths of a percent. We added the zero release at zero time since this is a physically reasonable boundary condition for this system. The 0.26 factor multiplying time in the exponent in Figure 3 was obtained from the fit in Figure 5.

**Figure 5.**
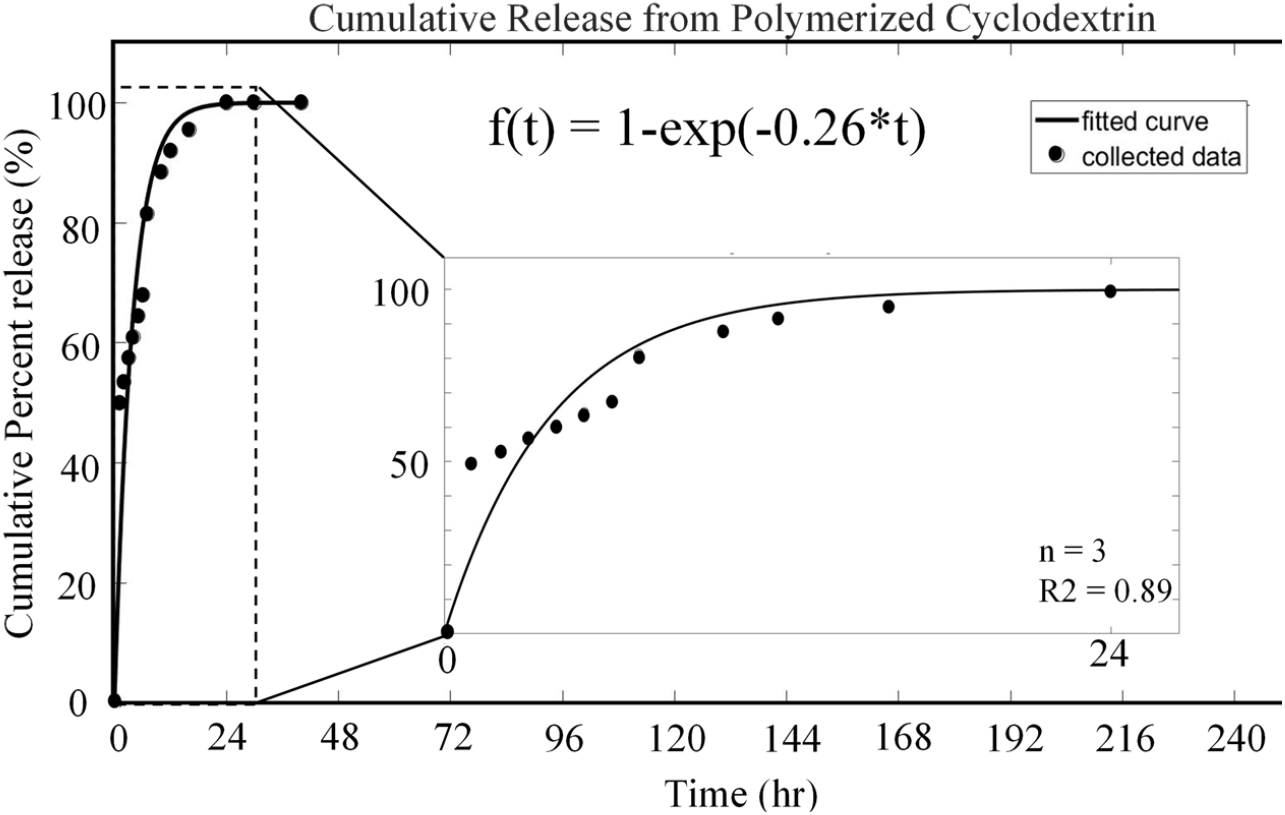
The cumulative release of doxycycline from polymerized cyclodextrin is plotted. Aliquots were taken every hour for 6 hours and then every day for 2 days and plotted as circles, with these time intervals being marked along the x-axis. The release amount is plotted as a percent of final cumulative amount released along the y-axis. The plot increases rapidly initially and plateaus around the 24 hour mark. After the plateau the plot increases very minimally, staying close to the asymptote of 1. A solid line is fitted to the data points. A model was created by fitting a curve in the form of y(t)=1-exp(Bcoll*t). The fitted equation is 1-exp(−1.26*t).

### Mathematical Modeling

Exponential models of the form shown in Eq. 1 were fit to the cumulative release plots. Polymerized cyclodextrin and collagen hydrogels were fit to Equation 1 with A_coll_ and A_CD_ set to zero. B_CD_=0.26 (Fig. 4) and B_coll_=1.31 (Fig. 5) were obtained from the fits. The two exponents were then used to fit Equation 1 to the cumulative release plot of doxycycline from cyclodextrin loaded collagen hydrogel: f(t) = 1 – A_CD_*exp(−0.26*t) – A_coll_*exp(−1.34*t). The “A” coefficients represent relative amounts of doxycycline released from the cyclodextrin and collagen portions of the gel. As shown in Figure 3, the best fit occurred with A_CD_=0.56 and A_coll_=0.43, R^2^=0.99.

## Discussion

### Incorporation of cyclodextrin oligomers into collagen hydrogels increased releasable drug compared to pure collagen hydrogels

Pure collagen gels released 1.6 mg of doxycycline whereas gels containing cyclodextrin released 3.6 mg of doxycycline, a 220% increase in releasable drug. Since variation in the masses of pure collagen gels was around 20%, we conclude the 220% increase was not due to variability in the size of the collagen gels. Instead, it is reasonable to assume that incorporation of cyclodextrin was responsible for this increase.

### The collagen/cyclodextrin system provides controlled long-term release of doxycycline

Cumulative release plots shown in figures 3-5 show an initial burst release within 24 hours followed by sustained release for 11 days consistent with published release studies from collagen (Semyari et al., 2018; Tihan et al., 2019) and cyclodextrin (Khodaverdi et al., 2021; Rivera-Delgado et al., 2016). In addition to cyclodextrin, doxycycline release can be manipulated by crosslinking collagen (Tihan et al., 2019). Because we crosslinked to maintain sample shape during drug elution, we thought it prudent to investigate the magnitude of this effect in our system. We modeled Tihan’s results using Eq. 1 with A_CD_=0. Their uncrosslinked group had a release exponent of B_coll_=2.5. Their most crosslinked group resulted in B_coll_=0.82, a 300% slower release than the uncrosslinked group. We saw a 500% difference in release rate between pure cyclodextrin polymers and pure collagen samples suggesting incorporation of cyclodextrin has a larger impact on release rate than just crosslinking. Since all our samples were similarly crosslinked, we do not expect differences between groups were a function of crosslinking.

Tihan’s work crosslinked with glutaraldehyde while we used genipin. Both act on terminal amines primarily on lysine and hydroxylysine in collagen molecules leading us to expect drug elution properties due to crosslinking by these agents to be roughly comparable. While not proven, the arguments above suggest the release rate of collagen gels can be tuned up to a factor of eight by utilizing both cyclodextrin oligomer incorporation and degree of crosslinking.

### Gels containing cyclodextrin oligomers show slower release rate than pure collagen hydrogels

Release trends of our experimental groups are qualitatively similar but not identical. We quantified these differences using the fitted exponential model. Insoluble polymerized cyclodextrin and pure collagen gels each have one mechanism of release. Therefore, the plots were fit with a single-exponential model: f(t) = 1 – exp(−B*t)(Albu et al., 2009). The resultant exponents, B_CD_ = 0.26 and B_coll_ = 1.31, show substantially slower release from cyclodextrin than from collagen alone, producing a more gradual approach towards the final release asymptote in Fig. 5 than in Fig. 4.

Gels containing cyclodextrin have both mechanisms of release: collagen and cyclodextrin. Therefore, the plot was fit with a double-exponential model: f(t) = 1 – A_CD_ *exp(−0.26*t) – A_coll_ *exp(−1.31*t). The coefficients in the arguments of the exponents were fixed based on the single-exponential fits. The “A” coefficients were determined by fitting and describe the relative amounts of drug released from cyclodextrin and collagen respectively: A_CD_ = 0.56 and A_coll_ = 0.43. 56% of doxycycline release came from cyclodextrin, 43% from collagen. The fitted coefficients are consistent with a majority of the release coming from the slower releasing cyclodextrin component allowing for a more controlled and long-term release profile.

### Proposed mechanism by which incorporation of cyclodextrin oligomers increases drug loading and decreases release rate in collagen gels

Bonfield discusses a cycle of ligand docking and dissociation for diffusion based drug release from insoluble polymerized cyclodextrin (Bonfield et al., 2024). In a system of neighboring cyclodextrin rings, a drug molecule could dock, dissociate, then diffuse until docking with another cyclodextrin in the network. Repeated cycles provide a more controlled and long-term release compared to diffusion alone. Pure cyclodextrin materials achieve this affinity binding coupled with frustrated diffusion via insoluble, high-molecular weight polymers of cyclodextrin. In unpublished work, we attempted to combine these large insoluble cyclodextrin polymers with collagen gels but found the cyclodextrin precipitated out of solution during fibrillogenesis preventing an intimate mixture of collagen fibrils and insoluble cyclodextrin polymer.

We then turned to incorporation of small oligomers into our gels. Previous work (Ding et al., 2023;) using FTIR showed hydroxyls from cyclodextrin hydrogen bonding with carbonyls from collagen. Quartz crystal microbalance studies confirmed binding of cyclodextrin with collagen (Majumdar et al., 2018) suggesting βCD binds with hydrophobic amino acids on collagen consistent with cyclodextrin monomers and oligomers becoming bound within a collagen network. If hydrophobic interactions dominated the CD/collagen association, then monomers might have their hydrophobic drug binding pockets sterically hindered by their initial binding to collagen.

Based on this argument, we tried larger oligomers settling on one with an average degree of polymerization of 66. While further work is necessary to prove it, our results are consistent with these oligomers being distributed throughout a network of fluid filled voids within the collagen gels. One or a few cyclodextrin oligomer hydrophobic regions may bind to a collagen molecule leaving several sites in each oligomer free to bind with doxycycline. The doxycycline could then bind, release and diffuse multiple times fairly rapidly from one site to another along a single oligomer and then more slowly between oligomers within the gel. This tortuous diffusional path may explain the slower release rate of our combined collagen/cyclodextrin material compared to collagen alone.

Unlike our results, Grier did not find improved drug release when incorporating β-cyclodextrin monomers into collagen-GAG scaffolds (Grier et al., 2018). Three differences between our systems may explain the different findings. First, our scaffolds are made solely from reconstituted type I collagen fibrils whereas Grier created composite scaffolds with type I collagen and chondroitin sulfate, a glycosaminoglycan. Second, we used doxycycline while Grier used TGF-β1 and BMP-2. Third, Grier incorporated monomers of βCD while we incorporated oligomers of γCD. Differences between the seven unit βCD and eight unit γCD exist but are not suspected to be major causes of our different outcomes. Instead, the monomer/oligomer difference between our two systems is a more likely explanation of our different outcomes. Single monomer interactions may directly occupy the hydrophobic binding region or block it through steric hindrance limiting the availability of cyclodextrin for drug binding and release. Oligomers have greater mobility reducing steric hinderance to provide a network of binding regions.

Recent work by von Recum and others shows affinity-based drug delivery systems having slower more sustained drug delivery than delivery based on diffusion alone and also that these polymers could be refilled in vivo following implantation(Cyphert et al., 2021; Young et al., 2022). Such in vivo “refilling” could then provide additional windows of drug release using the same device. The oral environment considered above could be an ideal setting for such a refiling scenario. An antibiotic mouth wash could be used to bathe the collagen–cyclodextrin hydrogel in new drug solution allowing it to refill with antibiotics, followed by additional windows of drug release. This could be extended as long as needed, perhaps for the life of the patient.

## Conclusions

Short soluble oligomers of γ-cyclodextrin incorporated into type I collagen hydrogels provide enhanced drug loading and control over release of doxycycline. The amount of releasable doxycycline was increased 220% compared to collagen hydrogels alone. The release rate was slowed by a factor of five as determined by mathematical modeling using a sum of two exponentials. Slightly more than half the releasable doxycycline in the combined collagen/cyclodextrin system came from the incorporated cyclodextrin. These findings may enhance future treatment strategies as peri-implant diseases remain a question without a complete answer.

## Funding Acknowledgement

We thank the Veterans Adminstration Advanced Platform Technology Steven Garverick Award for generous support of this work.

None of the authors have a conflict of interest related to the work described here.

Eric Trout and Steven Eppell contributed to: conception and design of the experiments, acquistion, analysis and interpretation of the data, drafting and critically revising the manuscript.

Horst von Recum contributed to: conception and design of the experiments, analysis and interpretation of the data, drafting and critically revising the manuscript.

Leena Palomo contributed to: conception and design of the experiments, interpretation of the data, drafting and critically revising the manuscript.

All authors gave their final approval and agree to be accountable for all aspects of the work.

